# The H-NS regulator plays a role in the stress induced by carbapenemase expression in *Acinetobacter baumannii*

**DOI:** 10.1101/2020.05.18.103317

**Authors:** Fanny Huang, Noelle Fitchett, Chelsea Razo-Gutierrez, Casin Le, Grace Ra, Carolina Lopez, Lisandro J. Gonzalez, Rodrigo Sieira, Alejandro J. Vila, Robert A. Bonomo, Maria Soledad Ramirez

## Abstract

Disruption of the histone-like nucleoid structuring protein (H-NS) was shown to affect the ability for Gram-negative bacteria to regulate genes associated with virulence, persistence, stress response, quorum sensing, biosynthesis pathways and cell adhesion. Here, we used the expression of metallo-β-lactamases (MBLs) known to elicit envelope stress by the accumulation of toxic species in the periplasm to interrogate the role of H-NS in *Acinetobacter baumannii*, together with other stressors. Using a multidrug-resistant *A. baumannii*, we observed that H-NS plays a role in alleviating the stress triggered by MBL toxic precursors and counteract the effect of DNA-damaging agents, supporting its role in stress response.

**Importance:** Carbapenem-resistant *A. baumannii* (CRAB) is recognized as one of the most threatening gram-negative bacilli. H-NS is known to play a role in controlling the transcription of a variety of different genes, including those associated with stress response, persistence and virulence. In the present work, we uncovered a link between the role of H-NS in the *A. baumannii* stress response and its relationship with the envelope stress response and resistance to DNA-damaging agents. Overall, we posit a new role of H-NS, showing that H-NS serves to endure envelope stress that could also be a mechanism that alleviates the stress induced by MBL expression in *A. baumannii*. This could be an evolutionary advantage to further resist the action of carbapenems.

---

*Acinetobacter baumannii* is a nosocomial pathogen frequently resistant to multiple drugs and causes a wide variety of infections with associated high mortality rates. Carbapenem-resistant *A. baumannii* (CRAB) have frequently been reported among hospital bailouts (1). In addition, CDC’s 2019 Antibiotic Resistance Threats Report moved CRAB into the urgent threats category (2). The expression of carbapenemases is critical for this organism to thrive under the selection pressure of these antibiotics in clinical environments. Instead, under permissive conditions (absence of antibiotics), the expression of some metal-dependent carbapenemases compromises the fitness of *A. baumannii*, triggering different responses associated to envelope stress (8). Despite the increased knowledge gained in recent years regarding *A. baumannii’s* epidemiology, pathogenicity and antimicrobial resistance (3, 4), how this pathogen responds to stressful environments is still not completely understood.

H-NS is a histone-like nucleoid structuring protein that serves as a global repressor, and has been shown to be involved in stress response in Gram-negative bacilli, such as *Vibrio cholerae* and *Escherichia coli* (5, 6). H-NS is known to protect the bacteria from environmental stresses through regulation of transcription and translation of virulence genes, quorum osmolarity, stress etc. (7, 8).

In *A. baumannii* the disruption of H-NS was found to affect the ability of this bacterium to regulate genes associated with persistence and virulence (9). However, the role of H-NS in stress response in *A. baumannii* has not been addressed yet. Here, we aimed to test the role of H-NS in *A. baumannii* stress response and how this could be linked with the success of multi-drug resistance *A. baumannii* in the hospital environment. Recent studies have shown that the production of certain MBL carbapenemases exert an envelope stress in an *A*. *baumannii* laboratory strain, resulting in growth defects (10). In this way, to study the role of H-NS in overcoming different kinds of stress, we utilized and evaluated the expression of three MBLs: NDM-1, VIM-2 and SPM-1, as stressors in the periplasmic space of AB5075 strain, and different known DNA-damaging agents.

Lopez *et al*. have shown that the inefficient processing upon translocation of non-frequent carbapenemases in *A. baumannii*, such as VIM-2 and SPM-1, compromises the bacterial fitness by triggering an envelope stress (10). Instead, expression of NDM-1 (a common resistance determinant in *A. baumannii*) is coupled to efficient processing, without causing any stress (10). In this way, this system represents a unique model to test envelope stress response since this stress can be regulated by varying the expression levels of MBLs, which directly affect the accumulation of toxic species in the periplasmic space.

To study the possible role of H-NS in envelope stress relief to overcome the expression of NDM-1, VIM-2 and SPM-1, growth curves of AB5075 and AB5075 Δ-*hns* expressing the different MBLs were performed. The mutant strain did not show impaired growth neither with the empty vector nor when expressing NDM-1 compared to the wild type strain (Fig. 1A-C). In line with previous studies, the expression of VIM-2 or SPM-1 affected the growth of AB5075. This effect was more pronounced in a Δ-*hns* background, indicating that the lack of H-NS impairs the growth of strains expressing SPM-1 and VIM-2 (Fig. 1B-C).

**Figure 1.**
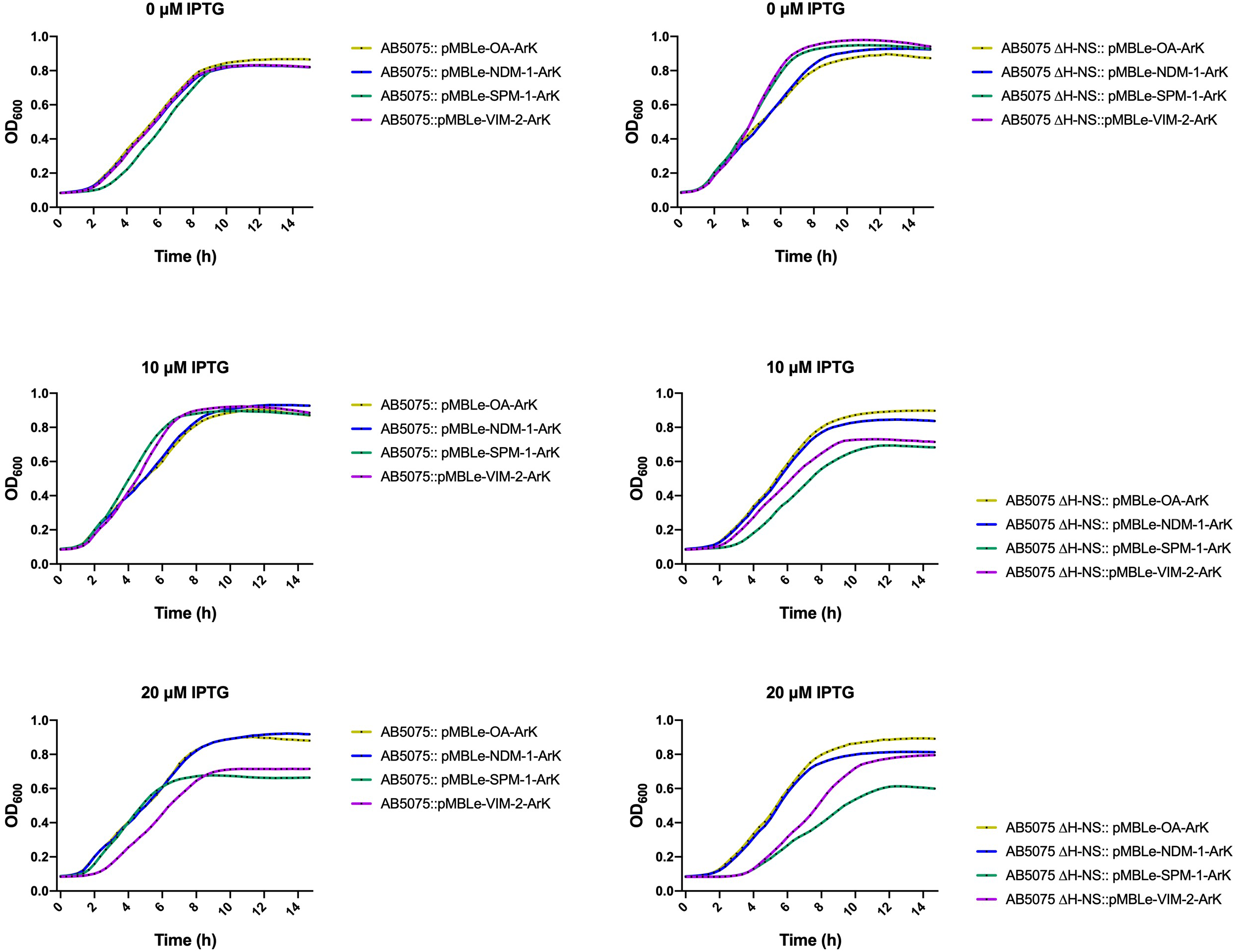
Growth curves of AB5075, and AB5075 Δ-*hns* strains, carrying the empty vector (pMBLe-OA) or expressing *bla*_NDM-1_, *bla*_VIM-2_, or *bla*_SPM-1_. Strains AB5075 and AB5075 Δ-*hns* with (pMBLe-OA-ArK, pMBLe-VIM-2-ArK, pMBLe-SPM-1-ArK, and pMBLe-NDM-2-ArK) were grown in LB broth plus A-B) 0, C-D) 10, or E-F) 20 µM IPTG. OD600 of the cultures was recorded every 20 minutes for 15 h. The data presented are the mean from 3 independent experiments.

Growth curves were unaltered when MBL expression was not induced (Fig. 1A-B) suggesting that H-NS plays a role in managing the accumulation of toxic precursor forms of SPM-1 and VIM-2. Our results also showed that when SPM-1 and VIM-2 were produced in relatively low amounts (0 and 10 μM IPTG), *A. baumannii* is able to withstand much of the impact on growth (Fig. 1A-D). The effect of fitness cost upon induction of SPM-1 and VIM-2 became evident at 20 μM IPTG (Fig. 1E-F).

We next sought to evaluate whether H-NS is also involved in the ability of *A. baumannii* to overcome other stressors, such as DNA-damaging agents MC and levofloxacin. AB5075 Δ-*hns* exhibited a decreased viability when exposed to MC (Fig. 2A). Also, the bacterial growth curve in the presence of levofloxacin showed an impaired growth for AB5075 Δ-*hns* (Fig. 2B). Overall, these data show that H-NS is involved in different *A. baumannii* stress responses.

**Figure 2.**
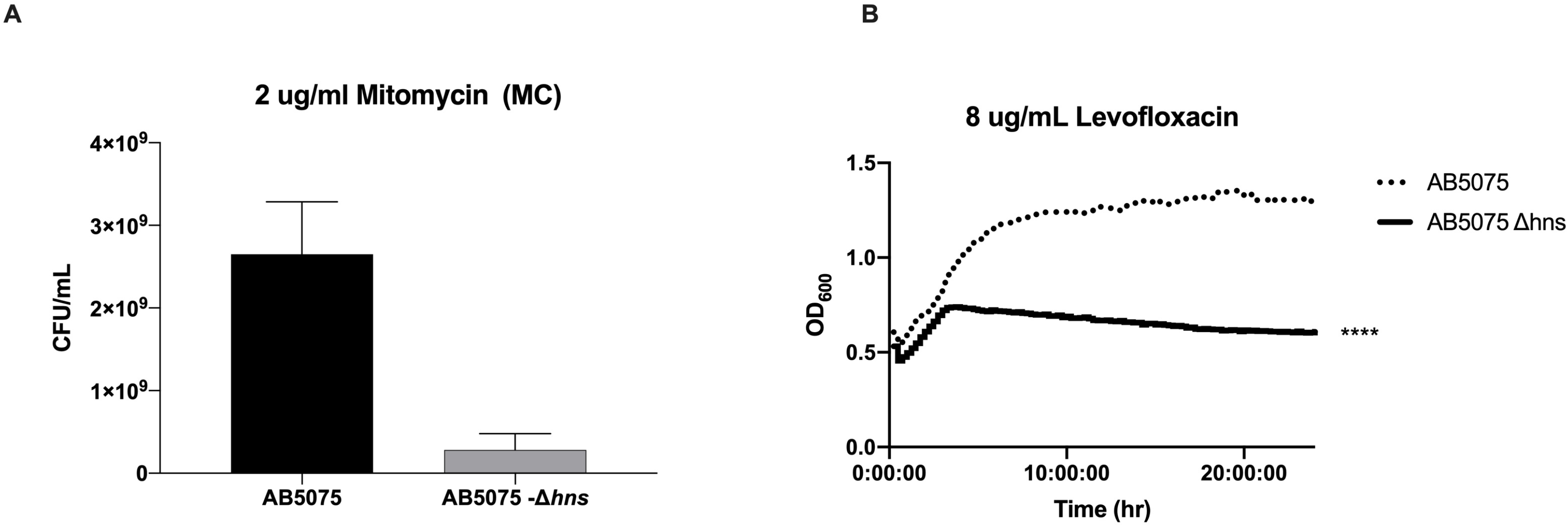
H-NS role to overcome DNA-damaging A) Mitomycin C (MC) survival assay of AB5075, and AB5075 Δ-*hns* strains. The cells were grown in LB broth overnight and then serially diluted in agar plates containing MC 0.2 ug/ml. The data presented are the mean +/− SD from 3 independent experiments. B) Growth curves of *A. baumannii* strains AB5075 and AB5075-*∆hns* in LB broth supplemented with 8 ug/mL levofloxacin. Growth was record (OD600) over 24 hours. Statistical analysis was performed using Mann-Whitney (n=3, *P-*value <0.05). The data presented are the mean from 3 independent experiments.

The stress response in *A. baumannii* is linked to limitation of essential nutrients, antibiotic treatment, oxidative damage, exposure to antiseptics, among others (11). When exposed to stress environments such as pleural fluid, *A. baumannii* can control the expression of different genes to overcome the stress and persist under the stressors signals (12).

In some gram-negative bacilli the role of H-NS in stress response has been well-characterized, e.g. in *V. cholerae* the deletion of *hns* has been shown to induce an envelope stress response causing an increasing expression of *rpoE*, and the regulators *rseA, rseB, rseC*, suggesting its role in cell envelope biogenesis (5). However, data on *A. baumannii* are scarce (9) (13).

Recent studies showed that periplasmic stress generated by production of toxic MBLs can be alleviated by an increase in the production of outer membrane vesicles (hypervesiculation phenotype) enclosing non-host-adapted MBLs. Along with membrane vesiculation, the activation of periplasmic proteases also acts to relieve the accumulation of toxic MBLs in the periplasm in non-frequent hosts (10). Here, we show a different strategy, involving the H-NS regulator, used by the highly resistant and hypervirulent AB5075 to cope with the expression of MBLs. We observed that AB5075 express NDM-1 without growth defects. Instead, the expression of VIM-2 and SPM-1 compromised *A. baumannii* survival, triggering a stress response H-NS-dependent.

We also observed that H-NS is involved in stress response, not only alleviating the stress imposed by expression of VIM-2 and SPM-1, but also by DNA damaging agents. The expression of SPM-1 in the mutant H-NS strain caused a more drastic decrease in growth compared to VIM-2.

Collectively, our observations suggest that H-NS serves to overcome envelope stress and could also be a possible mechanism that may allow to alleviate the stress induced by VIM-2 and SPM-1 in *A. baumannii*, further increasing its repertoire to resist the action of carbapenems.

## Bacterial strains and plasmids

AB5075 and AB5075 Δ-*hns* were used in the present study. For expressing the different *bla* genes (*bla*_VIM-2_, *bla*_SPM-1_ and *bla*_NDM-1_) in *A. baumannii*, plasmids constructions of the MBL variants already containing *bla*_VIM-2_, *bla*_SPM-1_ and *bla*_NDM-1,_ as well as the empty vector pMBLe-OA (10) were used as backbone to include the apramycin (ArK^R^) to generate the plasmids pMBLe-OA-ArK, pMBLe-VIM-2-ArK, pMBLe-SPM-1-ArK, and pMBLe-NDM-1-ArK to be used in the MDR strains AB5075 and AB5075 Δ-*hns.* MBL expression was induced with low concentrations of IPTG (10 and 20 μM), as indicated.

## Electroporation

Electro-competent *A. baumannii* AB5075 and AB5075 Δ-*hns* cells were prepared and mixed with 25 ng of plasmid DNA followed by electroporation with a Bio-Rad Gene Pulser instrument at 2.5 kV, 200 Ω, 25 µF. The electroporated cells were placed in recovery with 1ml of LB broth for 2 hours at 37 °C in a shaking incubator followed by culturing overnight at 37°C on LB agar containing 15 µg/ml apramycin. At least 10 colonies were picked to confirm the presence of the different plasmids. To confirm their presence, plasmid extraction followed by gel electrophoresis analysis and PCR reaction using the corresponding primers to amplify either *bla*_VIM-2_, *bla*_SPM-1_ and *bla*_NDM-1_, and ArK (apramycin resistant gene) were performed.

## Growth curves

Growth curves were conducted on 96-well plates in triplicate with strains AB5075, and AB5075 Δ-*hns* with (pMBLe-OA-ArK, pMBLe-VIM-2-ArK, pMBLe-SPM-1-ArK, and pMBLe-NDM-2-ArK) in LB plus 0, 10 or 20 µM IPTG. Overnight cultures were subcultured 1:50 in LB incubated for 15 hours at 37°C with medium shaking. Growth was measured at an OD_600_ every 20 minutes using a Synergy 2 multi-mode plate reader (BioTek, Winooski, VT, USA) and Gen5 microplate reader software (BioTek).

## DNA-damaging agents susceptibility assays

AB5075, and AB5075 Δ-*hns* cells were exposed to 0.2 μg/ml mitomycin C (MC) and cell count was performed to measure cell-killing as previously described (12). Assays were performed in triplicate, with at least three technical replicates per biological replicate. In addition, growth curve of strains AB5075, and AB5075 Δ-*hns* exposed to 0 or 8ug/ml of levofloxacin (sub-inhibitory concentration) were performed as described above measuring bacterial growth every 20 minutes using a Synergy 2 multi-mode plate reader (BioTek, Winooski, VT, USA) and Gen5 microplate reader software (BioTek).

## Transparency declarations

None to declare

## REFERENCES

1. Rodriguez CH, Nastro M, Famiglietti A. 2018. Carbapenemases in A*cinetobacter baumannii*. Review of their dissemination in Latin America. Rev Argent Microbiol 50:327–333.

2. CDC. 2019. Antibiotic Resistance Threats in the United States. Atlanta, GA: US Department of Health and Human Services, CDC; 2019.

3. CDC. 2019. Antibiotic resistance threats in the United States. Centers for Disease Control, Atlanta, GA.

4. Isler B, Doi Y, Bonomo RA, Paterson DL. 2019. New treatment options against carbapenem-resistant *Acinetobacter baumannii* infections. Antimicrob Agents Chemother 63.

5. Wang H, Ayala JC, Benitez JA, Silva AJ. 2015. RNA-seq analysis identifies new genes regulated by the histone-like nucleoid structuring protein (H-NS) affecting Vibrio cholerae virulence, stress response and chemotaxis. PLoS One 10:e0118295.

6. Dorman CJ. 2014. H-NS-like nucleoid-associated proteins, mobile genetic elements and horizontal gene transfer in bacteria. Plasmid 75:1–11.

7. Hurtado-Escobar GA, Grepinet O, Raymond P, Abed N, Velge P, Virlogeux-Payant I. 2019. H-NS is the major repressor of *Salmonella Typhimurium* Pef fimbriae expression. Virulence 10:849–867.

8. Ayala JC, Silva AJ, Benitez JA. 2017. H-NS: an overarching regulator of the *Vibrio cholerae* life cycle. Res Microbiol 168:16–25.

9. Eijkelkamp BA, Stroeher UH, Hassan KA, Elbourne LD, Paulsen IT, Brown MH. 2013. H-NS plays a role in expression of *Acinetobacter baumannii* virulence features. Infect Immun 81:2574–83.

10. Lopez C, Ayala JA, Bonomo RA, Gonzalez LJ, Vila AJ. 2019. Protein determinants of dissemination and host specificity of metallo-beta-lactamases. Nat Commun 10:3617.

11. Fiester SE, Actis LA. 2013. Stress responses in the opportunistic pathogen *Acinetobacter baumannii*. Future Microbiol 8:353–65.

12. Martinez J, Fernandez JS, Liu C, Hoard A, Mendoza A, Nakanouchi J, Rodman N, Courville R, Tuttobene MR, Lopez C, Gonzalez LJ, Shahrestani P, Papp-Wallace KM, Vila AJ, Tolmasky ME, Bonomo RA, Sieira R, Ramirez MS. 2019. Human pleural fluid triggers global changes in the transcriptional landscape of *Acinetobacter baumannii* as an adaptive response to stress. Sci Rep 9:17251.

13. Deveson Lucas D, Crane B, Wright A, Han ML, Moffatt J, Bulach D, Gladman SL, Powell D, Aranda J, Seemann T, Machado D, Pacheco T, Marques T, Viveiros M, Nation R, Li J, Harper M, Boyce JD. 2018. Emergence of High-Level Colistin Resistance in an *Acinetobacter baumannii* Clinical Isolate Mediated by Inactivation of the Global Regulator H-NS. Antimicrob Agents Chemother 62.

